# A *Drosophila* screen identifies a role for histone methylation in ER stress preconditioning

**DOI:** 10.1101/2023.03.10.532109

**Authors:** Katie G. Owings, Clement Y. Chow

## Abstract

Stress preconditioning occurs when transient, sublethal stress events impact an organism’s ability to counter future stresses. Although preconditioning effects are often noted in the literature, very little is known about the underlying mechanisms. To model preconditioning, we exposed a panel of genetically diverse *Drosophila melanogaster* to a sublethal heat shock and measured how well the flies survived subsequent exposure to endoplasmic reticulum (ER) stress. The impact of preconditioning varied with genetic background, ranging from dying half as fast to four and a half times faster with preconditioning compared to no preconditioning. Subsequent association and transcriptional analyses revealed that histone methylation, transcriptional regulation, and immune status are all candidate preconditioning modifier pathways. Strikingly, almost all subunits (7/8) in the Set1/COMPASS complex were identified as candidate modifiers of preconditioning. Functional analysis of *Set1* knockdown flies demonstrated that loss of *Set1* led to the transcriptional dysregulation of canonical ER stress genes during preconditioning. Based on these analyses, we propose a model of preconditioning in which Set1 helps to establish an interim transcriptional ‘memory’ of previous stress events, resulting in a preconditioned response to subsequent stress.

**Author Summary:** Stress preconditioning occurs when a history of previous stresses impacts an organism’s response to subsequent stresses. There are many documented cases of stress preconditioning, but the specific genes and pathways involved in the process are not well understood. Here, we take advantage of the natural genetic variation in the *Drosophila* Genetic Reference Panel to examine the role genetic variants play in modifying preconditioning outcomes. Our goal is to identify genes that contribute to the underlying mechanisms of preconditioning. Specifically, we measured preconditioning outcomes as the change in death rates of *Drosophila* on constant endoplasmic reticulum (ER) stress with and without heat stress preconditioning for each strain. We demonstrate that preconditioning outcomes are highly dependent on genetic background. Through association and transcriptional analyses, we found that histone methylation, transcriptional regulation, and immune status are all candidate pathways impacting preconditioning. Functional studies utilizing *Set1* knockdown flies demonstrated that Set1, a histone H3 lysine 4 (H3K4) methyltransferase enzyme, is critical for the proper expression of a subset of ER stress genes during preconditioning. Our data indicate that Set1 likely aids in creating a transient transcriptional ‘memory’ following initial stress that impacts the response to subsequent stress.

## Introduction

Organisms routinely face many stressors, including changes in temperature, viral infection, exposure to environmental toxins, hypoxia, and ischemia(1,2). Understanding how an individual can respond to numerous insults over a lifetime is an ongoing challenge. In laboratory environments, naïve cells and organisms are commonly used to dissect the mechanisms underlying stress response pathways. However, this does not reflect the complex history of stresses that would naturally occur.

Efficient responses to stresses are vital for producing and maintaining a healthy proteome. Therefore, cells have many canonical stress response pathways that combat cellular stresses. The endoplasmic reticulum (ER) is responsible for folding approximately 30% of all polypeptides, which is an error-prone process disrupted by many stressors(3,4). The ER stress pathway is one of the most thoroughly characterized canonical stress response pathways. Three ER membrane sensors, IRE1, ATF6, and PERK, detect misfolded proteins and respond by initiating the unfolded protein response (UPR). The UPR includes a robust transcriptional cascade that upregulates genes whose protein products refold or degrade misfolded proteins(3). If the cell cannot achieve ER homeostasis, then apoptosis occurs. An effective ER stress response is critical for healthy development and aging. Improper proteostasis and a decline in the ER stress response contribute to many diseases, such as Alzheimer’s disease, Parkinson’s disease, Type 2 diabetes, and more(5–9). An essential step in understanding how the ER stress response impacts disease is understanding how the stress response varies with genetic background and previous stress.

Natural genetic variation is a powerful tool for investigating canonical stress response pathways. Incorporating genetic variation into ER stress research has revealed new genes and pathways involved in the ER stress response(10–15). Much of what we understand about the ER stress response comes from studies that examine this response in isolation, using a single genetic background. In reality, ER stress occurs in a complex milieu of previous stresses that likely impact how the cell responds, and this likely varies with genetic background.

Preconditioning is a long-observed example of an organism’s ability to adapt to numerous assaults, whereby transient exposure to stress affects the organism’s ability to respond to subsequent stresses(16–19). There are many documented cases of preconditioning modifying the outcome of canonical stress pathways. For example, renal epithelial cells preconditioned with various pharmaceutical inducers of ER stress are resistant to subsequent peroxide-induced cell death(18). Exposing *C. elegans* to mitochondrial stress during larval development increases their ability to respond to and recover from heat stress as adults(20). In a mouse model of Parkinson’s disease, preconditioning mice with ER stress through tunicamycin injections is neuroprotective from subsequent 6-OHDA injections(16). Although this long-observed phenomenon has been documented, the molecular mechanisms are unknown, and stress response pathways continue to be studied in isolation.

We used natural genetic variation to understand the impact of stress preconditioning on the ER stress response. Here, we report the results of a stress preconditioning screen performed using a genetically diverse panel of *Drosophila melanogaster*. The screen revealed that preconditioning outcomes are strongly dependent on genetic background. We identified candidate modifier genes from association analyses of the preconditioning screen and predictive genes of preconditioning outcomes from transcriptional analyses. Together, these analyses revealed that immunity, transcriptional regulation, and histone methylation might play a role in the underlying mechanisms of preconditioning. Strikingly, we identified nearly all the components of the Set1/COMPASS complex as candidate modifiers. We investigated the impact of the loss of *Set1*, a conserved histone H3 lysine 4 (H3K4) methyltransferase in the Set1/COMPASS complex, on preconditioning. We found that *Set1* knockdown modifies preconditioning outcomes and leads to the transcriptional dysregulation of a subset of genes during preconditioning. We posit that after initial stress, Set1 plays a critical role in creating a transient transcriptional ‘memory’ of the event, resulting in a preconditioned response to subsequent stress.

## Results

### Genetic background modifies preconditioning outcomes

We subjected 177 strains of the *Drosophila* Genetic Reference Panel (DGRP) to a stress preconditioning screen to assess the impact of genetic variation on how preconditioning affects the ER stress response. The DGRP, a collection of fully-sequenced, inbred *Drosophila* strains derived from a natural population, is a powerful tool for uncovering novel effects of genotypic background on biological processes(21). For our preconditioning screen, we exposed males of each DGRP strain to heat stress preconditioning (or no preconditioning control), allowed them to recover for four hours, placed them on food containing tunicamycin (TM) to induce ER stress, and monitored survival (Fig 1A). Sublethal heat stress was chosen as the preconditioning stress because it is applied to flies uniformly, causes a robust response, has been extensively characterized, and begins within minutes of heat application(22–24). TM induces ER stress by inhibiting N-linked glycosylation and is commonly utilized in ER stress studies(11,25–28). TM-induced ER stress ultimately leads to the death of all flies(11). Therefore, the phenotypic outcome measured in this screen was the survival time under TM-induced ER stress.

**Figure 1:**
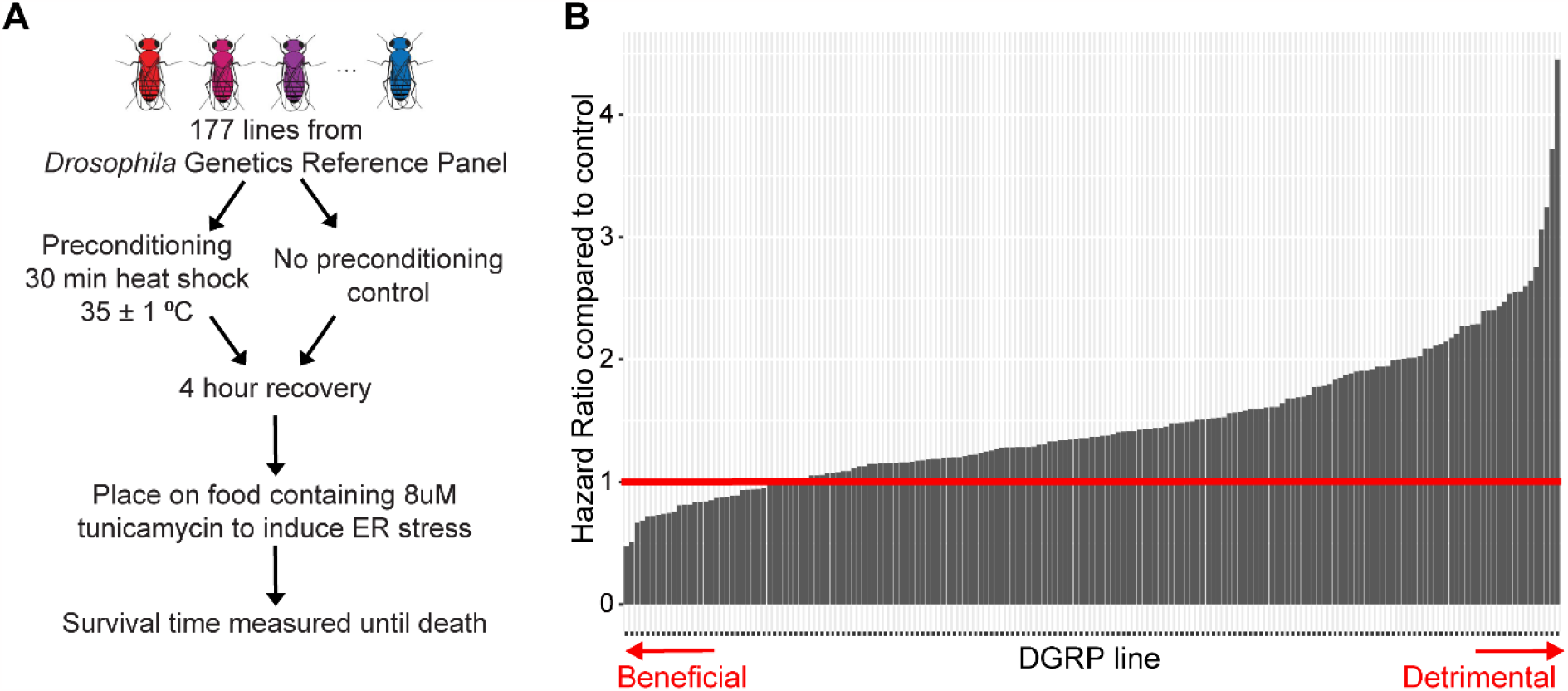
Genetic background alters the effect of preconditioning on ER stress survival times. (A) Experimental design of the stress preconditioning screen performed on 177 lines of the *Drosophila* Genetics Reference Panel (DGRP). (B) Results of stress preconditioning screen. Each bar represents a different DGRP strain. Cox proportional hazard ratio: [rate of death on ER stress with preconditioning]/[rate of death on ER stress without preconditioning]. A hazard ratio < 1 indicates preconditioning had a beneficial effect on ER stress survival, a hazard ratio = 1 indicates no effect, and a hazard ratio > 1 indicates a detrimental effect. The red line marks this transition point at hazard ratio = 1.

We compared survival curves of preconditioned and control flies from each DGRP strain to determine how preconditioning affects each strain’s response to ER stress. Survival was analyzed with the Cox proportional-hazards model to generate a hazard ratio for each strain that compares the death rate on ER stress with and without heat stress preconditioning(29) (S1 Dataset). A hazard ratio value of 1 indicates no significant change due to preconditioning, <1 indicates slower death in the preconditioned group compared to control (beneficial), and >1 indicates faster death in the preconditioned group (detrimental).

The stress preconditioning screen revealed that the effect of preconditioning on the response to ER stress is dependent on genetic background (Fig 1B). The impact of preconditioning on ER stress-induced death rates ranges from dying half as fast (hazard ratio = 0.47, p = 2.36 × 10^−7^) to 4.5 times faster (hazard ratio = 4.45, p < 2.0 × 10^−16^) compared to no preconditioning. There is no correlation in the DGRP between responses to preconditioning and previously reported ER stress responses (r = -0.16; p = 0.13)(11), heat tolerance (r = -0.047; p = 0.67)(30), or longevity (r = 0.036; p= 0.65)(31), indicating that these individual factors do not drive preconditioning outcomes (S1 Figure). Therefore, the variability in preconditioning outcomes directly results from unique, underlying genetic variation in the DGRP and is not simply the sum of the effects of variation on heat stress, ER stress, and overall longevity.

### Genome-wide association (GWA) analysis identifies candidate modifier genes of preconditioning

We performed a GWA analysis using the hazard ratios generated for each DGRP strain to identify candidate modifier genes underlying the variable preconditioning outcomes. We applied a linear mixed model to query 2,007,145 polymorphisms (MAF ≥ 0.05) to identify variants that are significantly associated with outcomes from our preconditioning screen (S1 Table).

Evaluating the role of any specific SNP is difficult due to limitations imposed by multiple testing. Therefore, in this study, we put little emphasis on individual polymorphisms. Instead, we prioritized identifying potential modifier genes, which has been a very successful approach in previous DGRP screens(11,12,14,32). If a SNP fell within an annotated gene, we assigned it to that gene. If a SNP was in an intergenic region, we assigned it to the closest gene within 1 kb. We did not evaluate SNPs that were more than 1 kb from the closest gene.

With an arbitrary p-value cut-off of p ≤ 1×10^−4^, the GWA analysis identified 110 polymorphisms associated with stress preconditioning. Of these 110 polymorphisms, 81 fell within 1 kb of a known gene (S2 Table). These 81 variants included seven downstream and four upstream of a gene, three in a 3’ UTR, one in a 5’ UTR, 56 in an intron, and nine in protein-coding exons. Of the nine in protein-coding exons, one is a non-synonymous variant (*CG15784*), and eight are synonymous.

These 81 polymorphisms are associated with 40 unique *Drosophila* genes (Table 1). Of these 40 genes, 22 have a human orthologue with a DIOPT score of at least five(33). The 40 genes do not include any canonical ER stress or heat shock response genes, reinforcing the observation that differences in response to these individual stresses do not drive the variation in preconditioning. Gene Ontology (GO) analysis did not identify any significant enrichment.

**Table 1:**
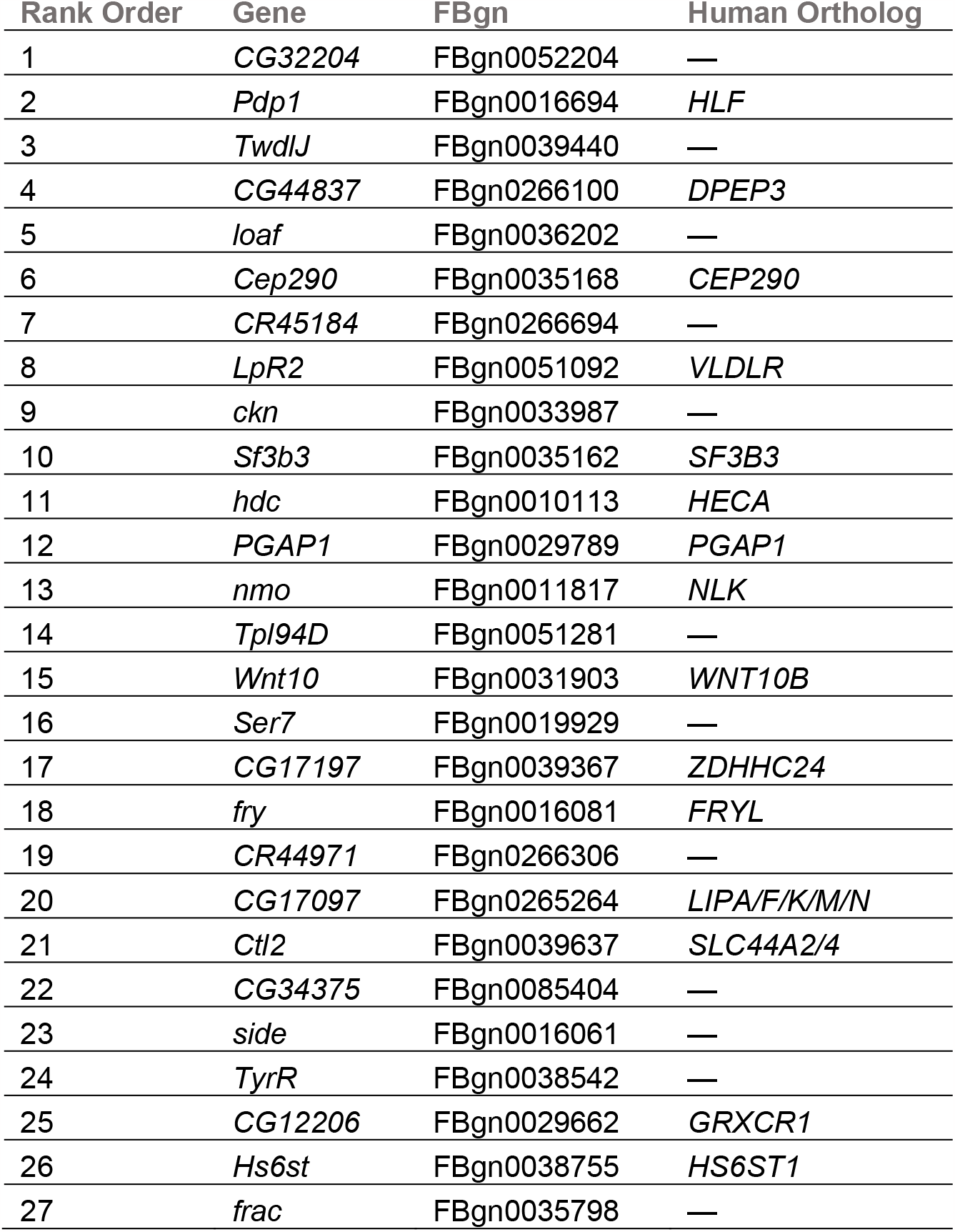
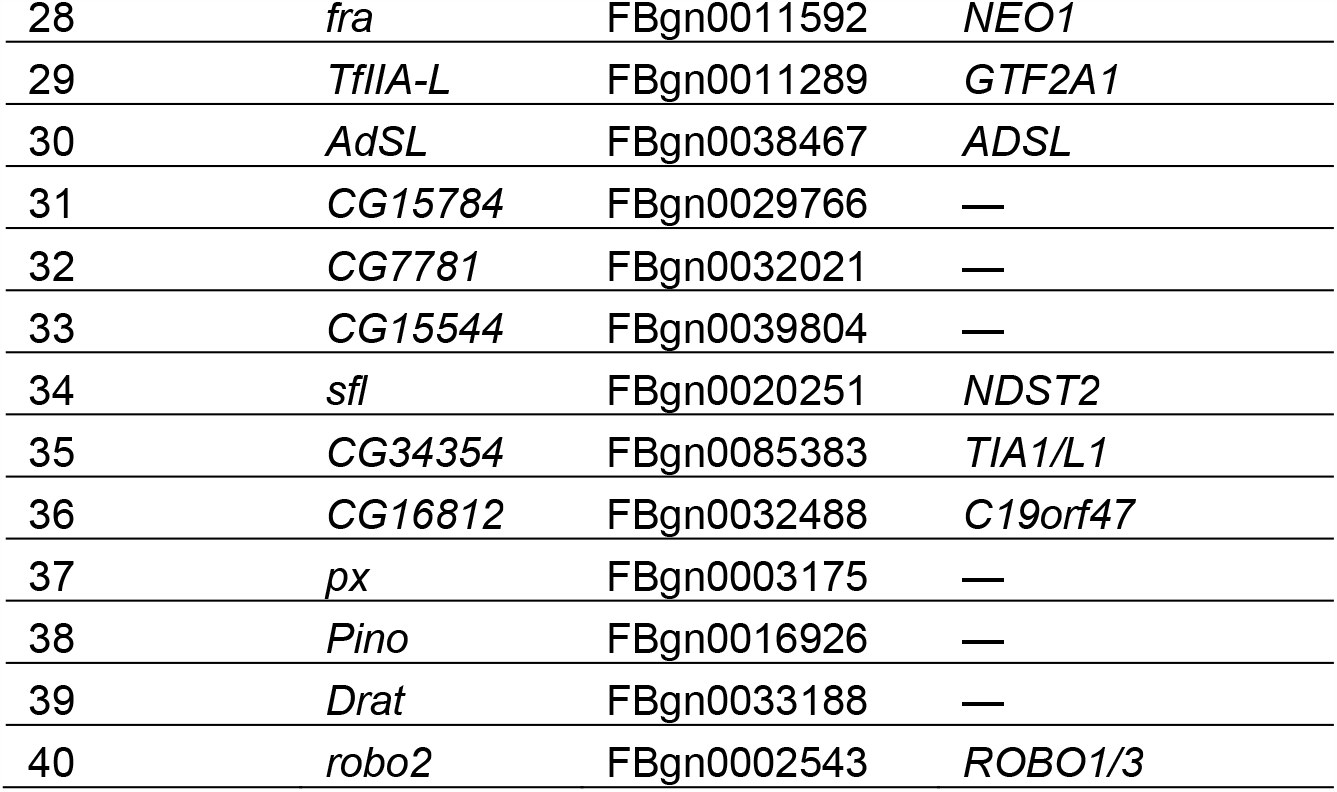
Candidate modifier genes identified from GWA. Modifiers are ranked based on the most significant SNP associated with each gene. Human orthologs include the ortholog with the greatest DIOPT score (≥5).

A subset of candidate modifiers identified in our GWA analysis indicated the potential involvement of chromatin organization and transcriptional regulation in preconditioning mechanisms. The GWA analysis identified *Tpl94D* (no human ortholog), an HMG-box domain protein with known roles in chromatin reorganization, indicating a potential role of chromatin organization in preconditioning outcomes(34). Candidate modifiers also include two general transcriptional factors, *Pdp1* and *TfIIA-L* (human orthologs: *HLF* and *GTF2A1*, respectively). *Pdp1 (HLF)* is particularly interesting because it is the top candidate modifier with a human orthologue. *Pdp1 (HLF)* is a widely expressed transcription factor critical for proper development and circadian rhythm(35,36). *TfIIA-L (GTF2A1)* is a member of the preinitiation complex required for RNA polymerase II initiation(37). The precise mechanism of how *Pdp1 (HLF)* and *TfIIA-L (GTF2A1)* impact preconditioning is unclear, but their known roles suggest the possible importance of transcriptional regulation in preconditioning.

### Gene set enrichment analysis (GSEA) uncovers the importance of histone methylation in preconditioning

Thus far, our investigation has only examined the candidate modifiers harboring individual polymorphisms that exceed a statistical threshold. Although this is a valuable method for identifying SNP-ranked candidate genes, it neglects most of the association data generated by our GWA analysis. In a second approach to analyzing the GWA results, we utilized Gene Set Enrichment Analysis (GSEA) to explore the entire association dataset (S1 Dataset). GSEA assigns each variant to a gene and calculates a gene-based p-value enrichment score that determines the significance of all variants assigned to the gene(14,32,38,39). The genes are then re-ranked by this new gene-based p-value. GSEA examines this newly ranked list and identifies GO terms enriched at the top of the list.

There were 11 enriched ontology terms uncovered by GSEA (Fig 2A; S3 Table) (p ≤ 0.05, enrichment score ≥ 0.50, number of genes contributing to ontology ≥ 5). Three gene ontology terms pointed to histone methylation as a key process regulating preconditioning (Fig 2A, indicated terms highlighted). Each of the three histone methylation ontology terms mostly contain unique genes with minimal overlap (Fig 2B).

**Figure 2:**
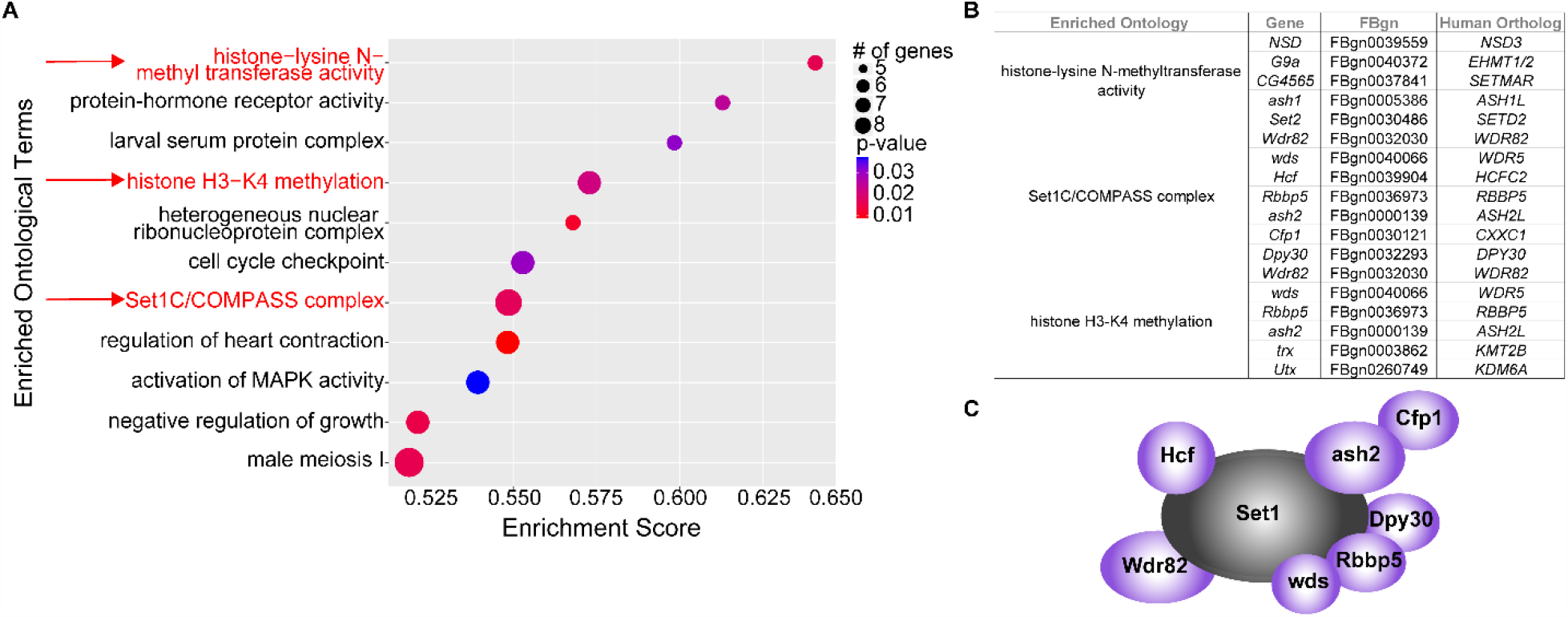
Gene set enrichment analysis (GSEA) reveals a role of histone methylation in preconditioning. (A) Top GSEA results. Ontology terms are listed by descending enrichment score. p-values are illustrated by the red to blue color gradient. The number of genes contributing to each category is indicated by the size of the circle. Ontology terms related to histone methylation are shown in red and indicated by a red arrow. Cutoffs include p-value ≤ 0.05, number of genes ≥ 5, and enrichment score ≥ 0.50. (B) Histone methylation ontology terms indicated in red are expanded to show the genes contributing to each ontology. FBgn and human ortholog (DIOPT score ≥ 5) are listed for each gene. (C) The entire Set1/COMPASS complex is illustrated with all known subunits. All subunits with polymorphisms contributing to the ontology terms related to histone methylation are shown in purple.

The most highly enriched ontological term is “histone-lysine N-methyltransferase activity” (GO:0018024). Of particular interest, the genes contributing to this enrichment include all the known *Drosophila* H3K36 methyltransferases, *NSD (NSD1-3), ash1 (ASH1L*), and *Set2 (SETD2*)(40–42) (Fig. 2B). These enzymes play roles in various processes, including transcription initiation and repression. Set2 (SETD2) associates with RNA polymerase II and plays a role in transcription elongation(43).

Another enriched ontological term from our GSEA is “Set1C/COMPASS complex” (GO:0048188). Strikingly, the genes contributing to the enrichment of this category include all subunits of the *Drosophila* Set1 complex, except Set1 itself(44) (Fig 2B-C). Identifying nearly every subunit of the Set1 complex demonstrates that genetic variation in each component contributes to the observed variation in preconditioning outcomes, illustrating the importance of the Set1 complex in preconditioning. Set1 (SETD1A/B) is responsible for the majority of histone H3K4 trimethylation in *Drosophila*(45,46). H3K4me3 marks histones proximal to the promoters of actively transcribed genes and promotes efficient transcription initiation through interaction with RNA polymerase II(45).

The final enriched ontological term associated with methylation is “histone H3-K4 methylation” (GO:0051568). This ontological term includes many genes associated with Set1 (SETD1A/B) (Fig 2B). Additionally, this ontology group includes *trx (KMT2A/B)*, another enzyme associated with H3K4 histone methylation in *Drosophila*(44–46). This category also includes *Utx (KDM6A)*, an H3K27me3 demethylase linked to transcriptional regulation that colocalizes with RNA polymerase II(47,48).

### RNAseq reveals potential gene expression predictors of preconditioning outcomes

Although GWA and GSEA uncovered a range of pathways that potentially underlie preconditioning mechanisms, the survival statistic is a culmination of many different processes. Expression differences in the basal state, heat shock response, recovery post-heat shock, and ER stress response might contribute to the ultimate survival outcome measured by the preconditioning screen. Thus, we sought to identify factors at early time points that might predict the outcome of preconditioning.

To identify genes with expression patterns predictive of preconditioning outcomes, we focused on the DGRP strains at the extreme ends of the distribution (Fig 1B). These strains included the five with the most beneficial preconditioning outcomes (RAL69, RAL93, RAL359, RAL387, RAL409) and the five with the most detrimental preconditioning outcomes (RAL195, RAL304, RAL335, RAL737, RAL819). We refer to these groups as the beneficial and detrimental groups, respectively. To identify a standard mechanism for detrimental or beneficial preconditioning outcomes, we treated the five strains in each group as technical replicates instead of investigating each strain individually. Combining five strains into a single group increases background noise introduced by genetic variation. Only genes with substantial, similar effects, irrespective of genetic background, will be detected as differentially expressed between the beneficial and detrimental groups.

RAL409 did not cluster with the other beneficial strains, as illustrated in PCA plots (S2 Figure A-C). Therefore, we removed RAL409 as an outlier for the following analyses. The PCA plots also reveal more variability in the detrimental group than in the beneficial group (S2 Figure C-D). This clustering pattern suggests that there is likely a common mechanism underlying a beneficial preconditioning outcome and that there may be more strain-specific mechanisms for a detrimental effect.

Under basal conditions, we found ten genes upregulated and seven genes downregulated in the beneficial group compared to the detrimental group (Fig 3A, Table 2) (padj ≤ 0.05, fold change ≥ 1.5). The two most highly upregulated genes in the beneficial group are *CG15263* (no human ortholog) and *CG6788* (Log_2_(fold change) = 8.267 and 4.042, respectively). With a DIOPT score of 4, *CG6788*’s predicted human orthologue is *FGL1*(33). While the function of *CG6788* is unknown, FGL1 is an immune checkpoint ligand that binds to LAG3, leading to T-cell depletion(49). *CG6788* may also play a role in immunity in *Drosophila*. The two most downregulated genes in the beneficial group, *Tep1 (CD109)* and *IM4* (no human ortholog)(Log_2_(fold change) = -1.860 and -1.086, respectively), participate in the *Drosophila* immune response by activating Toll(50,51). These results suggest that basal immune status may affect whether preconditioning is beneficial or detrimental.

**Figure 3:**
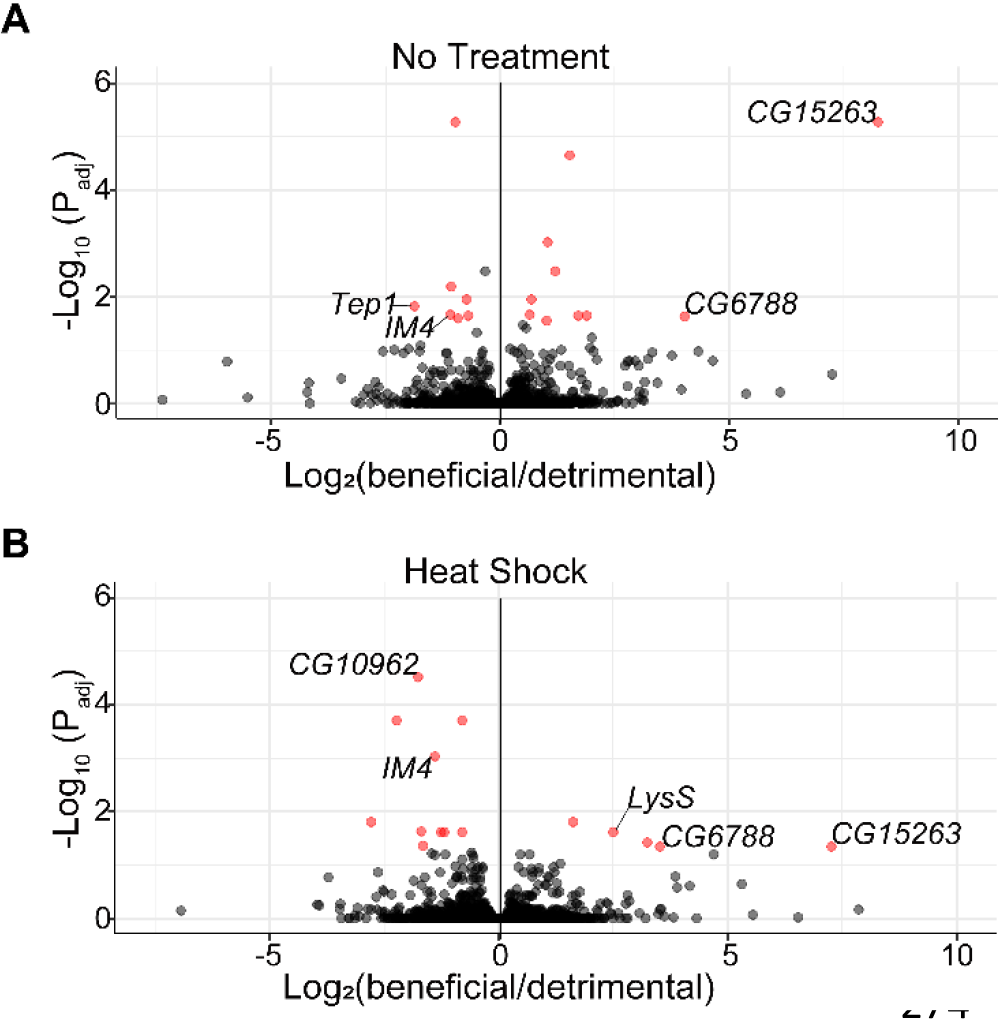
Differentially expressed genes in the beneficial group compared to the detrimental group. Volcano plots illustrating RNAseq results comparing beneficial/detrimental groups (A) with no stress or (B) immediately post-heat shock. Each point represents a different gene. Red points are significantly differentially expressed using significance cut-offs of padj ≤ 0.05 and fold change ≥1.5. Points to the right are upregulated in the beneficial group, and points to the left are downregulated. Labeled genes are discussed in the results.

**Table 2:**
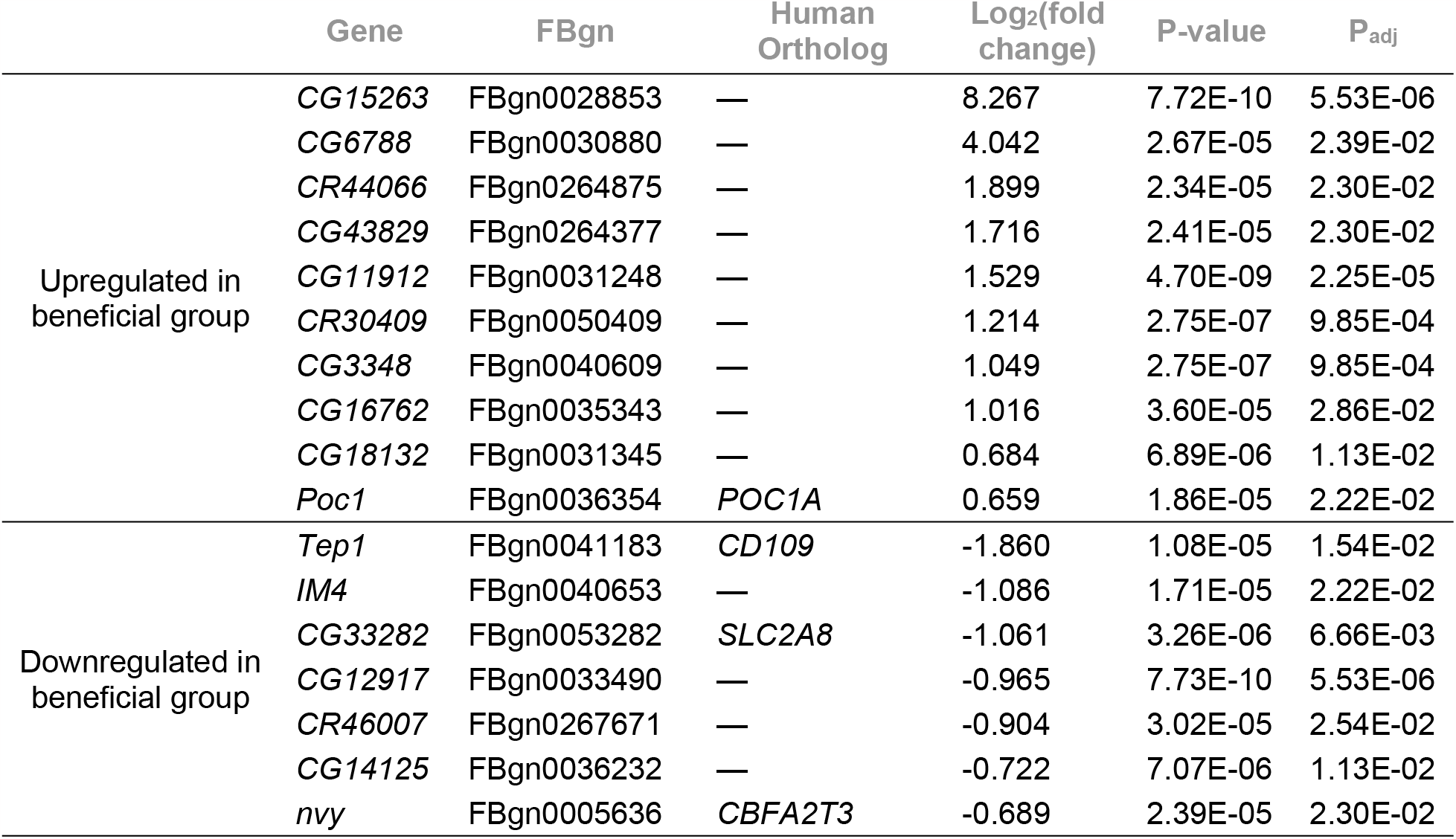
Differentially expressed genes in the beneficial group with no treatment. Upregulated or downregulated genes in the beneficial group compared to the detrimental group. Genes are ranked by Log_2_(fold change). Cut-offs include padj ≤ 0.05 and fold change ≥1.5. Human orthologs have a minimum DIOPT score of 5.

Post-heat shock, five genes are upregulated, and ten genes are downregulated in the beneficial group compared to the detrimental group (Fig 3B, Table 3) (padj ≤ 0.05, fold change ≥ 1.5). Two upregulated genes, *LysS (LYZ)* and *CG6788* (Log_2_(fold change) = 2.51 and 3.54, respectively), and one downregulated gene, *IM4* (Log_2_(fold change) = -1.37), are implicated in immunity(49,50,52). Of note, *CG15263* and *CG6788* remain the two most upregulated genes in the beneficial group, and *IM4* remains downregulated, regardless of treatment (no stress or post-heat stress). *CG15263* has no established function or human orthologue.

**Table 3:**
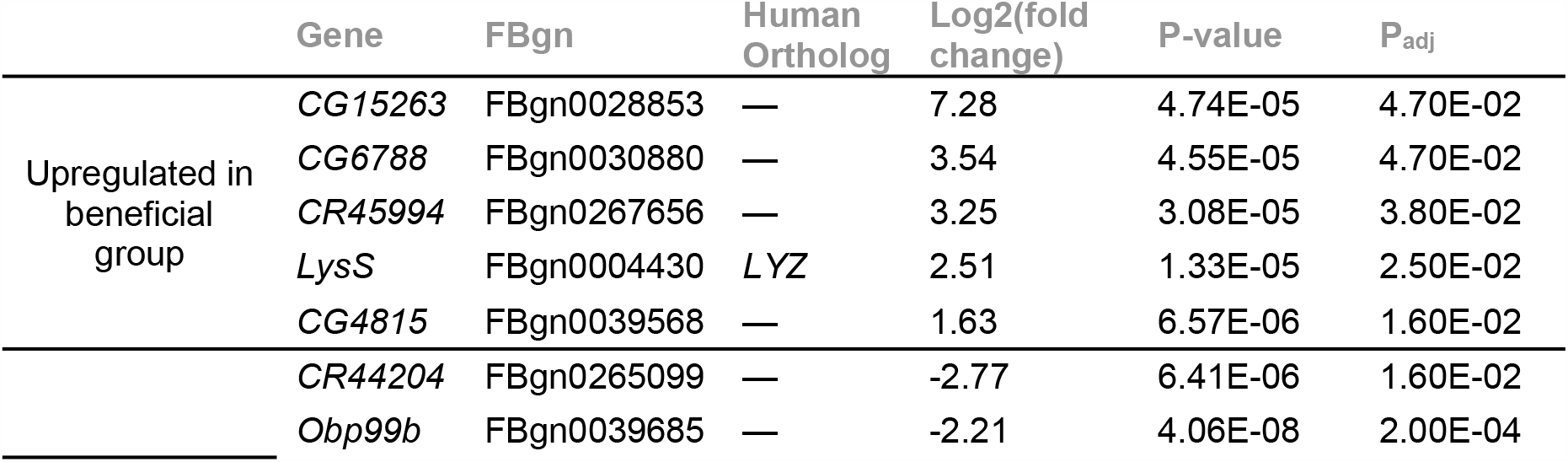
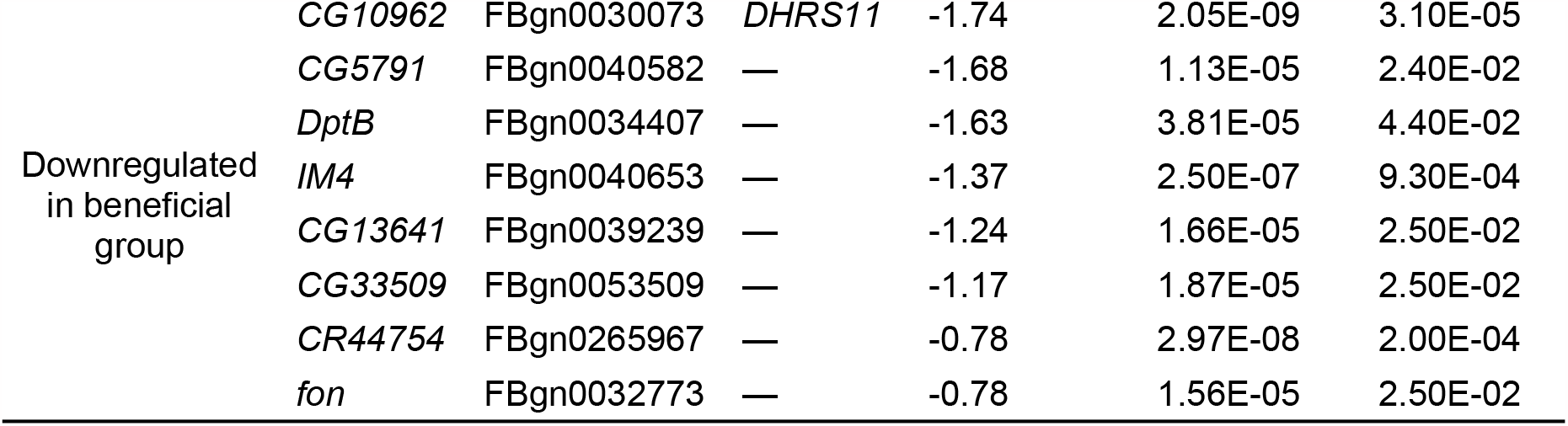
Differentially expressed genes in beneficial group post-heat shock. Upregulated or downregulated genes in the beneficial group compared to the detrimental group. Genes are ranked by Log_2_(fold change) within their group. Cut-offs include padj ≤ 0.05, and fold change ≥1.5. Human orthologs have a minimum DIOPT score of 5.

Another downregulated gene post-heat shock was *CG10962 (DHRS11)* (Log_2_(fold change) = -1.74), which may play a role in the ER stress response. A previous DGRP study found that a SNP in *CG10962 (DHRS11)* is strongly associated with changes in survival time in an environment of constant ER stress(11). *CG10962 (DHRS11)* is the only potential predictor of preconditioning outcomes associated with ER stress. We did not identify any significant ER stress or heat shock genes in this analysis, which reinforced the hypothesis that the individual stresses of preconditioning do not play a role in preconditioning mechanisms.

### Loss of *Set1* leads to increased variance in preconditioning outcomes

Our preconditioning screen and GWA analysis uncovered histone methylation and transcriptional regulation as candidate pathways for modifying responses to preconditioning (Table 1; Figure 2). We focused our initial functional investigation on *Set1 (SETD1A/B)* because all subunits bound to Set1 contain genetic variants contributing to preconditioning outcomes (Fig 2C). *Set1* is critical for the optimal transcription of active genes(45,53). Loss of *Set1* or its subunits leads to widespread impacts on gene expression(53). We knocked down *Set1* through ubiquitous expression of *Set1* RNAi (tubulin-GAL4 driver, “*Set1* KD”). This reduced *Set1* expression by approximately 55% (Fig 4A, S4 Table). This milder reduction in *Set1*, but not complete loss, better models the small effect sizes of natural variants identified in the preconditioning screen.

**Figure 4:**
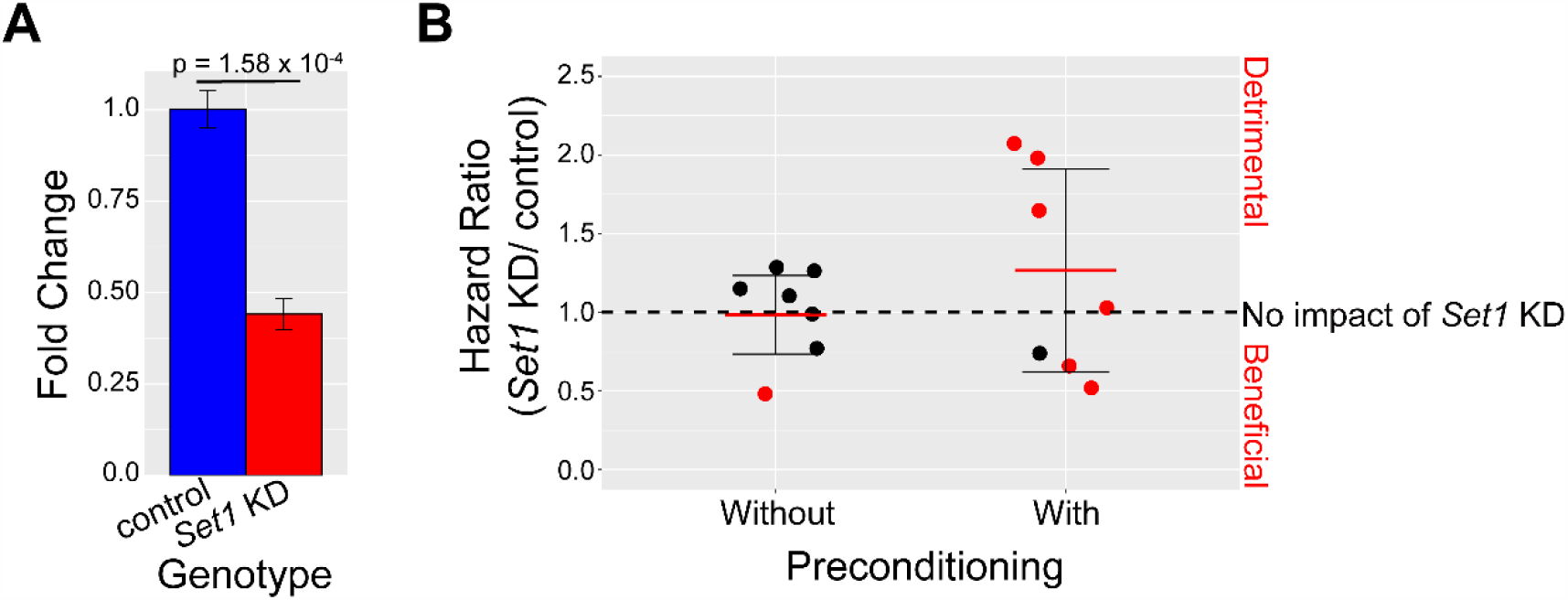
Impact of *Set1* KD on stress preconditioning. (A) *Set1* RNAi results in ∼55% reduction in *Set1* expression. (B) Plot of the hazard ratios comparing *Set1* KD to its genetically matched control without and with preconditioning. Each point represents the hazard ratio from a different replicate of 100 *Set1* KD and control flies. A hazard ratio < 1 indicates *Set1* KD had a beneficial effect compared to the control, a hazard ratio = 1 indicates no effect, and a hazard ratio > 1 indicates a detrimental effect. The dashed line indicates no effect (hazard ratio = 1) of *Set1* KD. The red line indicates the mean, and the cross bars indicate ±1 standard deviation. Red points indicate a statistically significant hazard ratio.

To investigate the impact of *Set1* KD on preconditioning, we utilized the same experimental design as the stress preconditioning screen (Fig 1A). In place of DGRP strains, we performed the assay using *Set1* KD and a genetically matched control. We utilized the Cox proportional-hazards model to generate a hazard ratio that compares *Set1* KD to the control with and without preconditioning (Fig 4B, S2 Dataset). We first tested whether *Set1* KD affects survival under ER stress without preconditioning. A hazard ratio of one indicates no significant change in ER stress survival without preconditioning between *Set1* KD and the control. Of seven replicates, only one replicate showed an effect (hazard ratio = 0.57, p-value ≤ 0.05) (Fig 4B). Therefore, *Set1* KD generally (6/7 replicates) does not impact ER stress survival times without preconditioning.

Next, we tested whether *Set1* KD affects survival under ER stress with preconditioning. More replicates show a significant effect due to the loss of *Set1* with preconditioning than without preconditioning. Four replicates result in a significant detrimental effect (hazard ratios= 2.14, 1.84, 1.76, 1.10, p-value ≤ 0.05), one results in no significant effect, and two replicates result in a significant beneficial effect (hazard ratios= 0.57, 0.57, p-value ≤ 0.05) of *Set1* KD on preconditioning outcomes. There was more variance in the impacts of *Set1* KD with preconditioning than without preconditioning (F test p-value = 0.037). These results indicate that *Set1* KD has a variable effect on preconditioning, but the phenotype we examined was not sufficiently precise to uncover the mechanism underlying this effect. We decided to investigate the impact of *Set1* KD at each individual step of the preconditioning assay in an effort to elucidate Set1’s role in preconditioning.

### Loss of *Set1* leads to dysregulation of critical stress response genes during preconditioned ER stress

ER stress and heat stress require robust transcriptional responses to refold misfolded proteins and return to homeostasis(11,54). *Set1* plays an active role in the efficient upregulation of stress response genes(45,53). The dysregulation of these transcriptional responses could lead to adverse effects on preconditioning. Therefore, we hypothesized that *Set1* KD alters the transcription levels of critical stress response genes, leading to abnormal mRNA levels at one or more steps of the preconditioning assay.

The stress preconditioning assay utilizes survival as a phenotypic readout. This provides insight into the cumulative effect of *Set1* KD on preconditioned ER stress, but does not reveal the individual contributions of each step of the preconditioning process (Fig 1A). Thus, we investigated the role of *Set1* in preconditioning at each step of the stress preconditioning assay, including without stress (no treatment), immediately following a 30-minute heat shock treatment (heat shock), immediately following a four-hour recovery from heat shock (post recovery), and immediately following a 16-hour TM treatment without and with preconditioning (ER stress and preconditioned ER stress, respectively).

We evaluated the role of *Set1* in upregulating canonical stress response genes at each stage of our preconditioning assay. We examined three established heat shock genes (*Hsp70, Hsp26, Hsp83*)(54,55) and three established ER stress genes (*Sil1, Ugt37A3, GstD2)*(11). We hypothesized that loss of *Set1* would lead to disrupted transcript levels of these critical stress response genes after one or more steps of the preconditioning assay, ultimately disrupting preconditioning outcomes. We first examined the heat shock and ER stress genes’ transcriptional responses to stress in control flies (S3 Figure). Next, we examined the transcriptional response of the same six genes in *Set1* KD flies exposed to the same preconditioning stress paradigm (S3 Figure). After examining how stress impacts these genes within each genotype, we analyzed the effect of genotype on transcriptional responses at each time point (control vs *Set1* KD) (Fig 5). This allowed us to measure how *Set1* KD altered gene expression throughout the preconditioning assay compared to the control.

**Figure 5:**
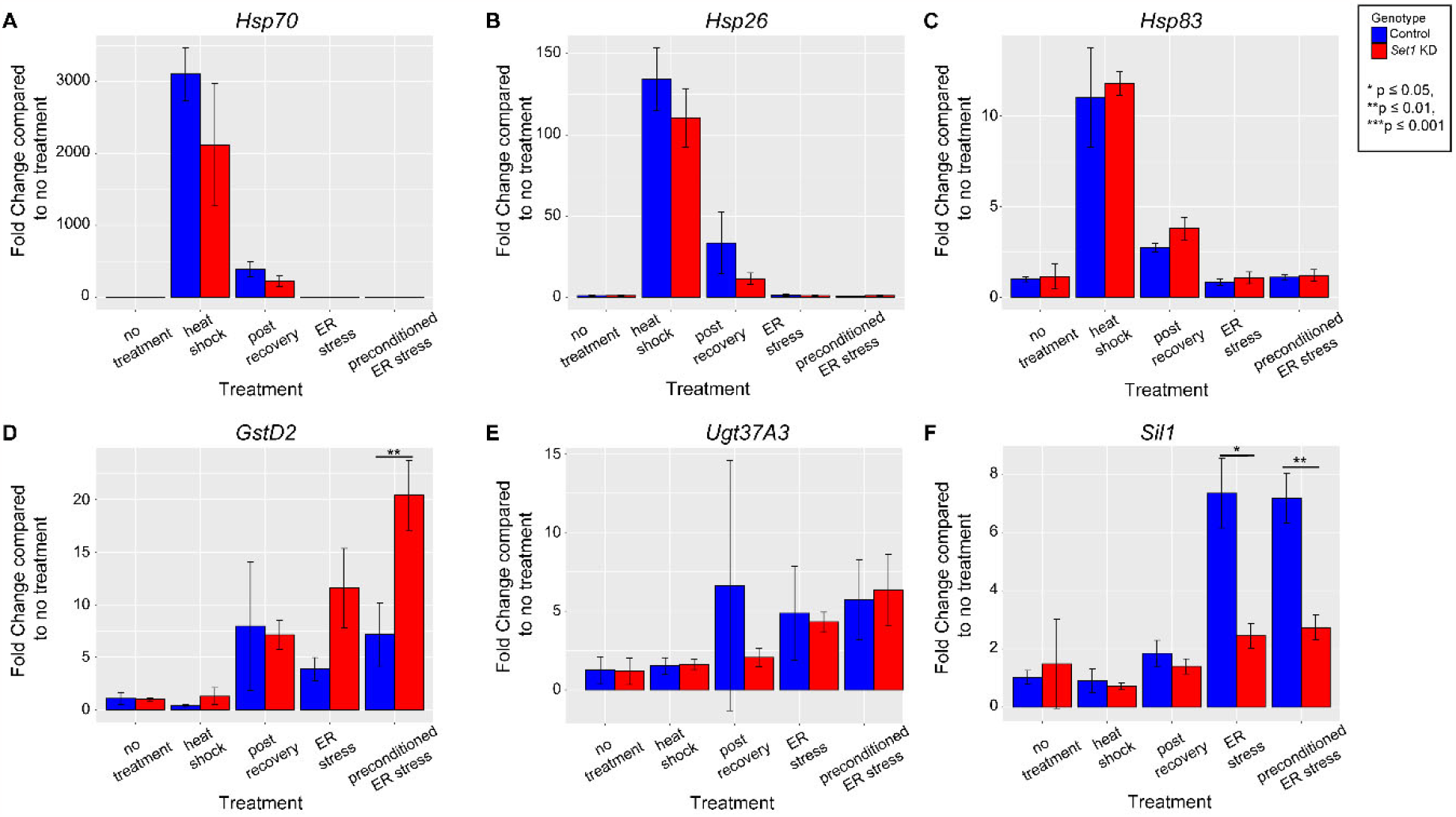
Set1 is necessary to regulate transcript levels of a subset of genes post-stress. Transcript expression change between controls (blue) and *Set1* KD (red). Paired t-test between control and *Set1* KD at each time point, significant differences are noted. (A-C) Heat stress genes. (D-F) ER stress genes.

None of the heat shock genes examined (*Hsp70, Hsp26, Hsp83*) were significantly impacted by *Set1* KD compared to the control at any point of the preconditioning assay (Fig 5A-C). They all showed a robust response to heat shock. This suggests that *Set1* is not necessary for normal heat shock gene expression during preconditioning. Similar to the heat shock genes, the ER stress gene *Ugt37A3* showed no significant impact of *Set1* knockdown across the five time points examined (Fig 5E).

In contrast, during preconditioned ER stress, *Set1* KD resulted in the misregulation of the ER stress genes *GstD2* and *Sil1. GstD2* and *Sil1* expression were not significantly altered by loss of *Set1* at any earlier timepoints. Following preconditioned ER stress, *GstD2* was upregulated in *Set1* KD flies more than in controls (Fig 5D; *Set1* KD: 20.42 ± 3.33; control: 7.17 ± 3.01-fold; p ≤ 0.001). *Sil1* displayed significantly lower expression in *Set1* KD flies compared to controls after ER stress without preconditioning (Fig 5F; *Set1* KD: 2.45 ± 0.43; control: 7.34 ± 1.19; p ≤ 0.05) and with preconditioning (Fig 5F; *Set1* KD: 2.73 ± 0.43; control: 7.18 ± 0.85; p ≤ 0.001). Taken together, our results suggest that *Set1* is necessary to regulate normal expression of a subset of critical ER stress response genes during preconditioning.

## Discussion

Organisms require an effective ER stress response for healthy development and aging. Disruption of the ER stress response underlies many human diseases – from diabetes to neurodegeneration(5–9). Therefore, a comprehensive understanding of the ER stress response’s fundamental biology is critical for informing future therapeutic development. The ER stress response naturally occurs within a complex history of other stresses that impact the response. Preconditioning is the phenomenon of transient exposure to stresses affecting responses to subsequent stresses. Research investigating the ER stress response has primarily focused on isolated stress events. To investigate how preconditioning impacts the canonical ER stress response, we utilized natural genetic variation in *Drosophila* to perform an unbiased screen.

We found that preconditioning outcomes vary greatly, depending on genetic background. Preconditioning outcomes in the literature primarily report a beneficial impact of preconditioning, but we found that this outcome is genotype-specific. Although several DGRP strains displayed a beneficial preconditioning outcome, many show neutral or detrimental effects. The spectrum of unique preconditioning outcomes in our screen made it possible to identify several candidate pathways that drive variation in outcomes. Discovering unexpected biological connections is a significant advantage of performing an unbiased genetic screen. First, our association analysis generated 40 candidate modifier genes of preconditioning. The list of candidate modifiers included two general transcriptional factors, *Pdp1 (HLF)* and *TfIIA-L (GTF2A1)*, and one gene involved in chromatin organization, *Tpl94D* (no human ortholog). Second, our GSEA identified multiple pathways that harbor genetic variants that impact preconditioning outcomes. The GSEA results included several ontologies involved in histone methylation. These results suggest a connection between transcriptional regulation, histone methylation, and preconditioning. Third, our RNAseq experiment uncovered 29 genes whose gene expression patterns may predict preconditioning outcomes. These different approaches did not identify any canonical ER stress or heat shock genes, which illustrates that the pathways underlying preconditioning are separate from the canonical pathways of the individual stressors.

Our unbiased approach revealed many intriguing new avenues for exploration. The association analyses identified two heparan sulfate sulfotransferases, *Hs6st* (*HS6ST1*) and *sfl* (*NDST2*). Heparan sulfate genes play a role in the cellular response to misfolded proteins(56). Candidate modifier genes also included two genes involved in lipid homeostasis, *LpR2* (*VLDLR*) and *CG17097* (*LIPA/F/K/M/N*). *LpR2* (*VLDLR*) is involved in neutral lipid uptake into cells, and *CG17097(LIPA/F/K/M/N*) is a triacylglycerol lipase involved in lipid metabolism and uptake(57,58). We previously demonstrated that disrupting cellular fatty acid composition through the disruption of *Baldspot/ELOVL6* impacts the ER stress response in *Drosophila*(13). The differentially expressed genes between beneficial and detrimental preconditioning groups included a previously unknown connection between preconditioning and immune pathways. Interactions between immune and stress responses have been previously observed(59). For example, immune peptides have been reported to block apoptosis signals triggered by stress response pathways(60).

Preconditioning has an established role in providing robust neuronal protection from subsequent brain injuries after initial stress(19,61). In one instance, a paradigm of heat stress protected rat brains from neuronal death by localized ischemia(62). Although many cases of preconditioning lead to neuroprotection, a genetic link between neurological function and preconditioning is unknown. Our association analysis identified several candidate modifier genes of preconditioning that encode proteins with known roles in axon guidance and neural development, including *loaf* (no human orthologue)(63), *ckn* (no human orthologue)(64), *sidestep* (no human orthologue)(65), *frac* (no human orthologue)(66), *fra* (*NEO1*)(67), and *robo2* (*ROBO1/3*)(68). Future studies are required to elucidate the underlying mechanisms of preconditioning in the brain.

We prioritized *Set1 (SETD1A/B*) for our initial functional validation because the association analysis and GSEA both indicated a potential role of chromatin organization and transcriptional regulation in preconditioning. Although the analyses identified several histone methylation genes, the Set1/COMPASS complex was the only candidate with genetic variants in all its subunits, except Set1 itself. The Set1/COMPASS complex marks promoter-proximal histones at actively transcribed genes. Set1 is a histone H3 lysine 4 (H3K4) methylase highly conserved from yeast to humans(46,55). Yeast, *Drosophila*, and humans have one, three, and six Set1/COMPASS family members, respectively(45,46). In *Drosophila, Set1* is responsible for the bulk of H3K4 di- and trimethylation but the other two Set1/COMPASS family members, Trr and Trx, contribute to di- and tri-methylation to a lesser extent(45). The redundancy between these family members may explain the variable effect we observed in the impact of *Set1* KD on stress preconditioning outcomes (Fig 4).

The ‘stress memory’ model of preconditioning proposes that cells acquire and retain epigenetic marks during initial stress, allowing them to ‘remember’ the stress and respond more efficiently to subsequent stress(69–71). This model provides a potential explanation of how Set1 is involved in preconditioning. In plants exposed to sequential droughts, transcript levels of a subset of stress-responsive genes are elevated more quickly during secondary dehydration, compared naive dehydration(72). After the drought state was resolved, these plants retained high levels of H3K4 trimethylation at the ‘primed’ stress-responsive genes, and RNA polymerase II also stalled at these genes(72). Yeast previously exposed to salt stress has 77 genes that activate more rapidly when exposed to subsequent oxidative stress(71). During stress in yeast, actively transcribed genes acquire H3K4 di- and trimethylation marks, and genes that acquire a ‘memory’ for stress retain H3K4 dimethylation after stress is resolved. Once again, these marks are associated with RNA polymerase II binding and retention at promoters. In vertebrates, the loss of Set1/COMPASS complex members leads to global misregulation of gene expression(73). In human HeLa cells, hundreds of genes show a more rapid transcriptional upregulation in response to IFN-γ after previous exposure to IFN-γ, and this ‘memory’ is associated with H3K4 dimethylation and poised RNA polymerase II at the promoters of these genes(74). Set1 is required for transcriptional memory in yeast and human cells.

In *Drosophila*, Set1 is actively recruited to stress response genes following stress application(45). Loss of *Set1* leads to less efficient upregulation of *Hsp70* and *hsp83*. Our study did not recapitulate these results, most likely due to differences in our experimental design. Kusch et al. utilized *Drosophila* S2 cell lines and larval salivary glands instead of whole adult flies and assayed more time points post heat stress. Kusch et al. reported H3K4 trimethylation marks promoter-proximal to *Hsp70* and *hsp83* were significantly reduced in *Set1* KD cells following heat shock, and RNA polymerase II displayed increased stalling at these promotors. Disruption of RNA polymerase II kinetics, such as in promoter-proximal pausing, is known to lead to transcriptional dysregulation(75). The role of H3K4 dimethylation post stress was not investigated in this study.

We established that disruption of *Set1* leads to dysregulation of a subset of stress response genes, particularly the ER stress genes. We propose that Set1 plays a role in preconditioning by establishing transcriptional ‘memory’ of stress events. We hypothesize that in *Drosophila*, Set1 adds H3K4 methylation marks (di- and tri-) promoter-proximal to stress-responsive genes during stress, and a subset of these marks are retained and influence how RNA polymerase II interacts with these genes to alter future transcriptional responses. To validate this model, further investigations into the global transcriptional and epigenetic impacts of *Set1* KD during preconditioning will need to be explored.

## Materials and methods

### *Drosophila* lines and maintenance

Flies were maintained at 25°C on a standard diet based on the Bloomington Stock Center standard medium with malt and without soy flour. Flies were on a 12-hour light/dark cycle. For the stress preconditioning screen, DGRP strains were obtained from the Bloomington *Drosophila* Stock Center. For the *Set1* functional work, a *Tubulin-*GAL4 driver (Bloomington *Drosophila* Stock Center: 5138), Attp40 (36304), and Set1 RNAi (40931) were used.

### Stress preconditioning assay

177 strains from the DGRP were used for the stress preconditioning screen. For each strain, 200 males were collected and placed in ten vials of 20 males each. Each male was between 2-8 days old and had recovered a minimum of two days since their last exposure to CO_2_ when they were exposed to stress. For preconditioning, five vials of 20 flies (100 flies total) were heat shocked by placing them into empty vials and submerging in a 35 ±1 °C water bath for 30 minutes. All heat shocks were performed in the morning, between 9 am and 12 pm. All flies were placed back on standard media and allowed to recover at 25 °C for 4 hours. TM food consisted of 8µM TM (Sigma-Aldrich CAS Number: 11089-65-9) dissolved in DMSO (Millipore Sigma CAS Number: 67-68-5), 1.3% agarose (BioRad #1620102), and 1% sucrose (Millipore Sigma CAS Number: 57-50-1) in DI water (similar to previous studies(11)). After recovery, all flies were transferred into vials containing 5 ml of TM food to induce ER stress. Flies were monitored every two hours during the light cycle (8am-8pm MST) and the number of dead flies were recorded. The control, no preconditioning flies, were treated in the same manner, but were not exposed to heat shock.

The *Set1* KD stress preconditioning assay used the same protocol. The only adjustment is that flies were monitored every two hours between 8am-12am MST (additional 4 hours during the dark cycle) once death was observed.

### Hazard Ratios

Survival analysis was performed using the Survival package in R (R version 4.2.0; survival package version 3.3-1; running under Windows 10 x64). The coxph test was performed to calculate the Cox proportional hazards ratio (HR) for each strain. The HR compares the death rate of the preconditioning group to the death rate of the control group within each strain. The HR takes into account all 100 flies exposed to heat stress and then ER stress and all 100 control flies exposed to only ER stress.

### Genome wide association (GWA) study

GWA was performed as previously described(12). DGRP genotype files were downloaded from the website: http://dgrp2.gnets.ncsu.edu/data.html. Variants were filtered for MAF (≥0.05) and non-biallelic sites were removed. The HR calculated from the preconditioning screen for 177 DGRP lines was regressed on each SNP. GEMMA (v. 0.94)(76) was used to estimate a centered genetic relatedness matrix and perform association tests using the following linear mixed model:

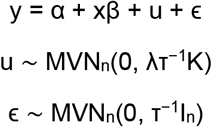

where, as described and adapted from Zhou and Stephens (58), y is the n-vector of the HR for the n lines, α is the intercept, x is the n-vector of marker genotypes, β is the effect size of the marker. u is an n x n matrix of random effects with a multivariate normal distribution (MVNn) that depends on λ, the ratio between the two variance components, T^−1^, the variance of residuals errors, and where the covariance matrix is informed by K, the calculated n × n marker-based relatedness matrix. K accounts for all pairwise non-random sharing of genetic material among lines. ϵ, is an n-vector of residual errors, with a multivariate normal distribution that depends on T^−1^ and I_n_, the identity matrix.

SNPs were assigned to genes within +/- 1 kb using the variant annotation file based on FB5.57 (dgrp.fb557.annot.txt) from the DGRP website, http://dgrp2.gnets.ncsu.edu/data.html. If multiple genes are within +/- 1 kb from a given SNP, the SNP was assigned to a single gene by prioritizing the variant type as follows: exon > UTR > intron > upstream or downstream. Human orthologues for each fly gene were chosen based on the greatest DIOPT score(33), with a minimum DIOPT score of 5.

### Gene Set Enrichment Analysis (GSEA)

GSEA analysis was performed as previously described (14,32,77). All polymorphisms from the stress preconditioning GWA (S1 Dataset) were assigned to a gene as described in the above section. Genes were organized into a rank-list based on their enrichment for polymorphisms and genes were assigned to GO categories. GSEA determines whether the top of the newly generated rank-list is enriched in genes belonging to any GO categories or if the genes in the category are randomly distributed throughout the list. Calculation of enrichment score was performed as described by Subramanian et al. 2005 (for code see Figshare: https://doi.org/10.25387/g3.9808379). Only GO categories with a corrected p-value ≤ 0.05, number of genes ≥ 5, and an enrichment score ≥ 0.50 were considered.

### RNAseq

mRNA sequencing was performed on total RNA from whole male 2-day old flies (10 flies per group sample). DGRP lines RAL69, RAL93, RAL359, RAL387, and RAL409 made up the beneficial group and RAL195, RAL304, RAL335, RAL737, and RAL819 made up the detrimental group. Flies were given 48 hours to recover from their last CO_2_ exposure. The no treatment flies were frozen down at the same time as the heat shock samples. The heat shock samples from each line were heat shocked for 30 minutes at 35 ±1 °C and then frozen down immediately post heat shock.

mRNA sequencing was performed on 20 samples (10 genotypes x 2 treatments x 1 replicate). RNA was extracted using a Direct-zol RNA Miniprep (Zymo Research R2061) using TRIzol Reagent (ThermoFisher Cat # 15596026) and including the DNAse step. Samples were prepared and sequenced by the Huntsman Cancer Institute High-Throughput Genomics Core. The 20 samples were sequenced on the NovaSeq 50 × 50 bp Sequencing, for a total of approximately 25 million paired reads per sample. Fastq files were trimmed using seqtk v1.2 software (for FastQ and processed files see GEO repository: GSE226958). RNAseq reads were aligned to the *Drosophila melanogaster* reference genome (assembly BDGP6.28, Ensembl release 102) using Bowtie2 v2.2.9 software(78) and alignment files were sorted and converted using Samtools v1.12(79).

Read counts were normalized using the default normalization method in DESeq2(80) package in R. Principle components analysis (PCA) was performed to identify outliers (S2 Figure). RAL409 was identified as an outlier and was removed for further analyses. The remaining samples were reanalyzed using Deseq2 v1.28.1. Differentially expressed genes in the beneficial group (compared to detrimental group) were identified before treatment and immediately post-heat shock. Remaining samples were renormalized and assessed using linear models with the DESeq2 package. Genes were considered significantly differentially expressed if the adjusted p-value ≤ 0.05 and the fold change magnitude was ≥ 1.5 (Log_2_ fold change ≥ 0.585 or ≤ -0.585).

### RT-qPCR

Each sample contained 12 adult male flies that were 4-7 days old and had been off CO_2_ for 3 days when exposed to stress. 30 samples were collected (5 timepoints x 2 genotypes x 3 replicates) and RNA was extracted using a Direct-zol RNA Miniprep (Zymo Research R2061) using TRIzol Reagent (ThermoFisher Cat # 15596026) and including the DNAse step. RNA was converted to cDNA using a ProtoScript® II First Strand cDNA Synthesis Kit (NEB Cat # E6560L). RT-qPCR was performed using a QuantStudio 3 96-well 0.2 ml block instrument and PowerUp SYBR Green Master Mix (ThermoFisher Cat # A25741). We used primers from the FlyPrimerBank located at http://www.flyrnai.org/flyprimerbank:

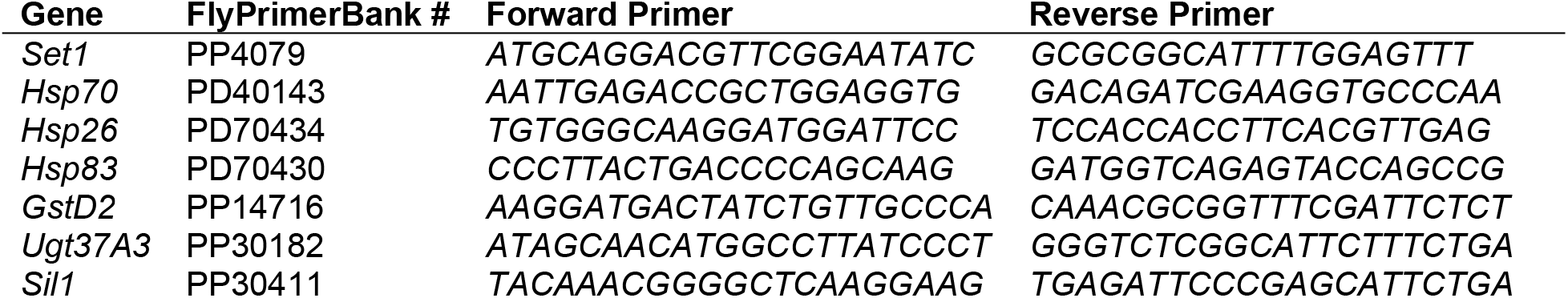

Results were analyzed using the Delta-Delta Ct method.

## Acknowledgments

Stocks obtained from the Bloomington *Drosophila* Stock Center (NIH P40OD018537) were used in this study. We thank Emily Coelho and Shayna Scott for technical assistance with fly management. KGO was supported through the NIH/NIGMS Genetics T32 Fellowship from the University of Utah (T32 GM007464). This research was supported by the NIH through an NIGMS R35 award (R35GM124780) (CYC).

## Supplemental Information

**S1 Dataset: Stress preconditioning screen results**. ER stress survival with and without preconditioning and hazard ratios are provided for each DGRP line, and tabs are grouped by strain. Each fly’s survival time on TM, treatment (preconditioning or control), and replicate group is provided. The first tab includes a summary of each strain’s hazard ratio, p-value, and ln(hazard ratio).

**S1 Figure: Correlation plots comparing stress preconditioning screen results to previously reported DGRP impacts on ER stress, heat tolerance, and longevity**. The distribution of stress preconditioning screen hazard ratios across DGRP strains is not correlated with (A) the distribution of the hazard ratio of death rates of 114 DGRP lines on TM-induced ER stress compared with drug-free control food(11); (B) the distribution of maximum heat tolerance of 100 DGRP lines(30); (C) the distribution of the mean lifespan of males across 186 DGRP lines at 25°C(31).

**S1 Table: Complete GWA results for stress preconditioning screen**. Alleles are listed with chromosome position, rs ID, and significance.

**S2 Table: Top SNPs from GWA**. The top 81 SNPs from the GWA analysis are listed with chromosome location, FBgn ID, associated gene, variant type, distance from gene, major and minor allele, allele frequency, and significance. Cutoffs include: SNPs +/- 1 kb of a known gene, af ≥ 0.05, p ≤ 0.0001.

**S3 Table: Top enriched GSEA categories**. GSEA categories are ranked in descending order by enrichment score. Genes with polymorphisms contributing to each ontology are indicated as FBgns. Cutoffs include p-value ≤ 0.05, number of genes > 4, and enrichment score ≥ 0.50.

**S2 Figure: PCA plots of RNAseq results**. The color of each point indicates the treatment group, and the shape indicates the preconditioning group. (A) PCA plot of RNAseq of five DGRP strains with beneficial preconditioning outcomes and five with detrimental outcomes without treatment. (B) PCA plot of RNAseq on the same strains as (A) immediately post-heat shock. (C) Heat shock versus no heat shock PCA plot for beneficial strains. (D) Heat shock versus no heat shock PCA plot for detrimental strains.

**S4 Table: qPCR results**. Reports dCt and 2^-ddCT values for the following target genes: *Hsp70, Hsp26, Hsp83, Sil1, Ugt37A3, GstD2*, and *Set1*. The genotypes queried include *Set1* KD and Attp40 control. Flies were collected without treatment, immediately after heat shock, after a four-hour recovery, and after a 16-hour TM treatment with and without preconditioning. *Set1* was only investigated at the no treatment timepoint to evaluate knockdown efficiency.

**S2 Dataset: Set1 KD stress preconditioning assay lifespan data**. The first tab includes a summary of each replicate’s hazard ratios (Set1 KD/ control with and without preconditioning), and p-value. The following tabs include the survival time on TM on hours, genotype, heat shock treatment, and vial for each fly in each replicate. Each tab is a different replicate.

**S3 Figure: Analysis of stress response gene expression post-stress relative to no treatment**. Each plot illustrates qPCR fold change data for one of six stress response genes in control (blue) or *Set1* KD (red) flies. (A-C) Genes upregulated immediately after heat stress. (D-F) Genes upregulated post-ER stress.

