## Supplementary figures and images for "A *Drosophila* screen identifies a role for histone methylation in ER stress preconditioning"

### Supplementary figure 1

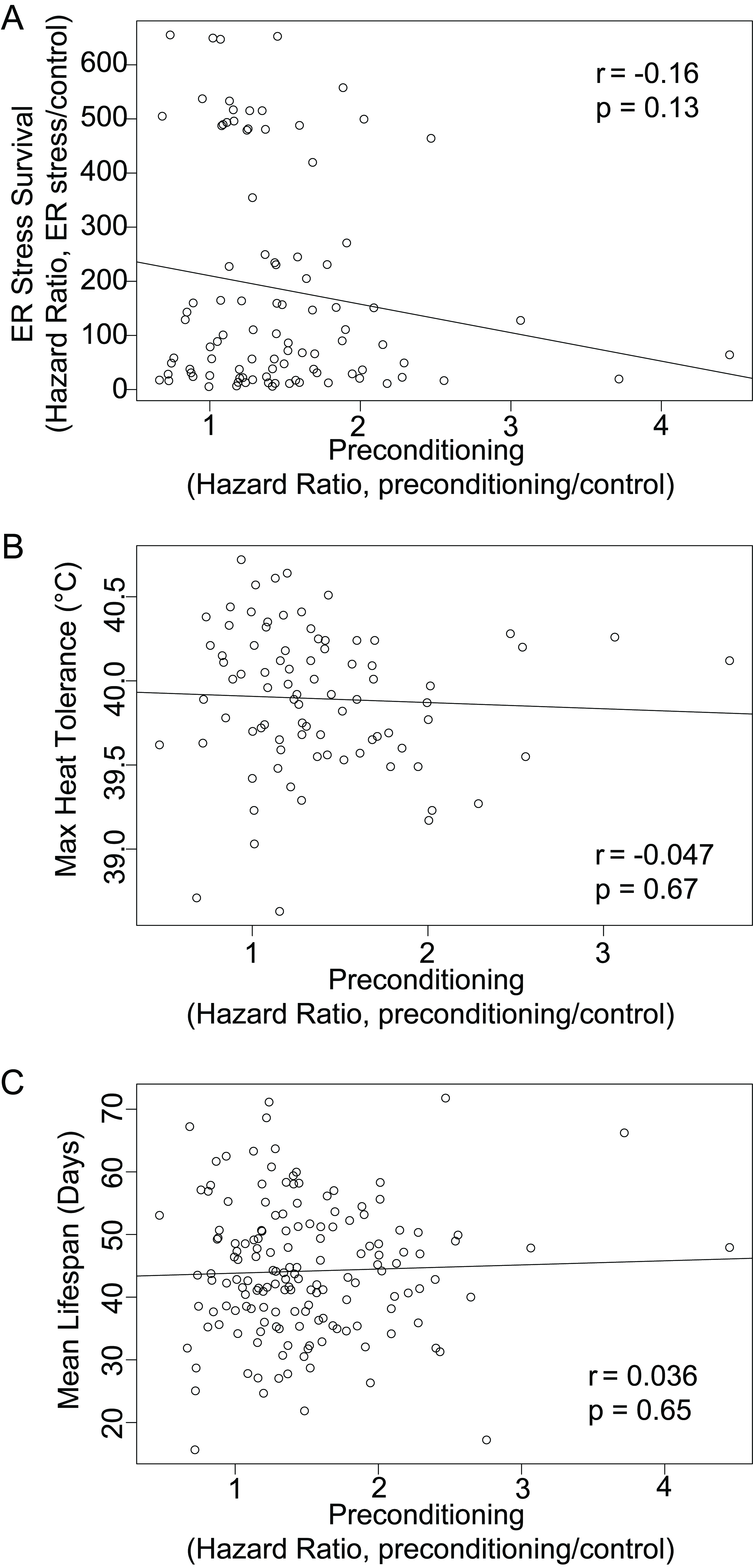

### Supplementary figure 2

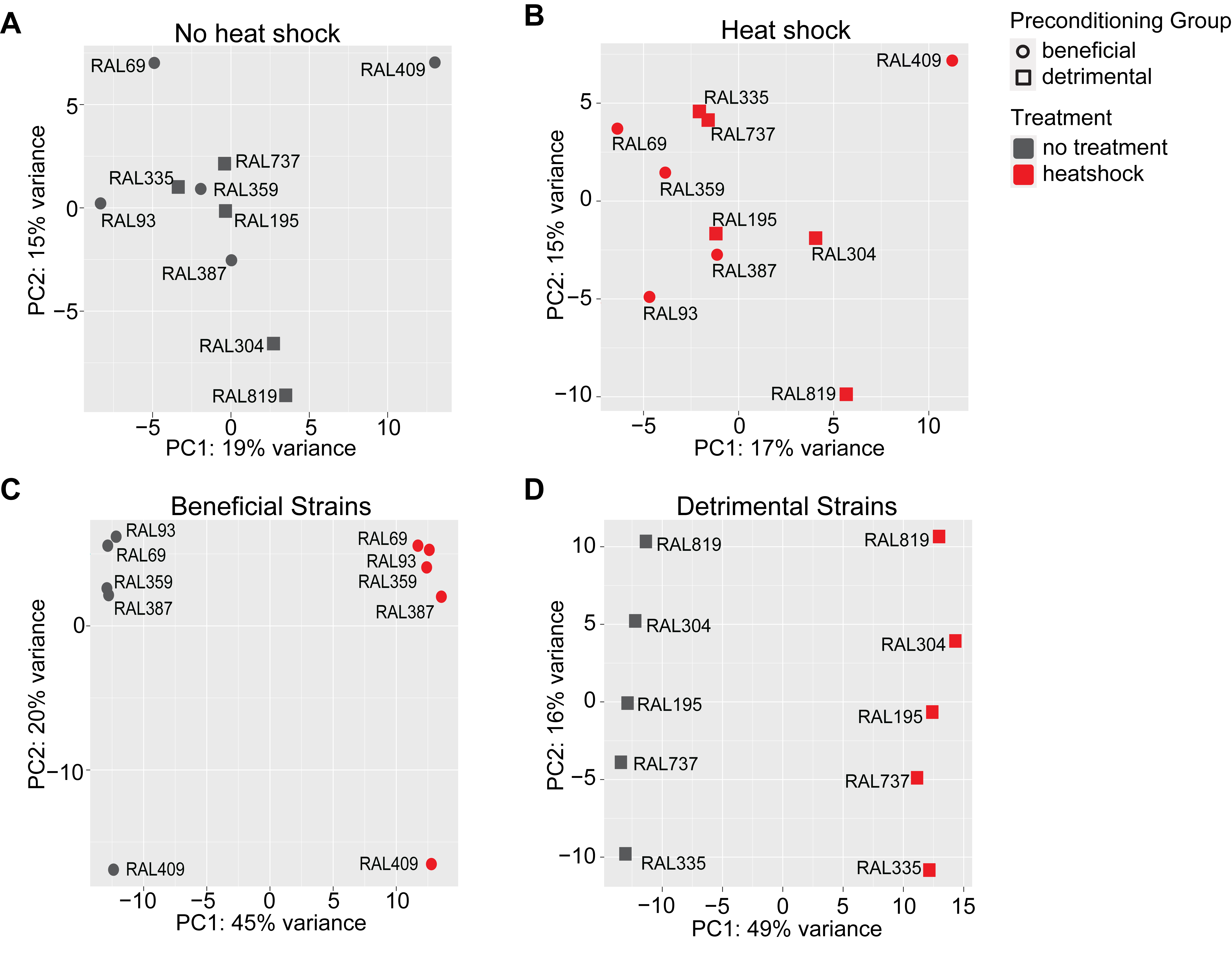

### Supplementary figure 3

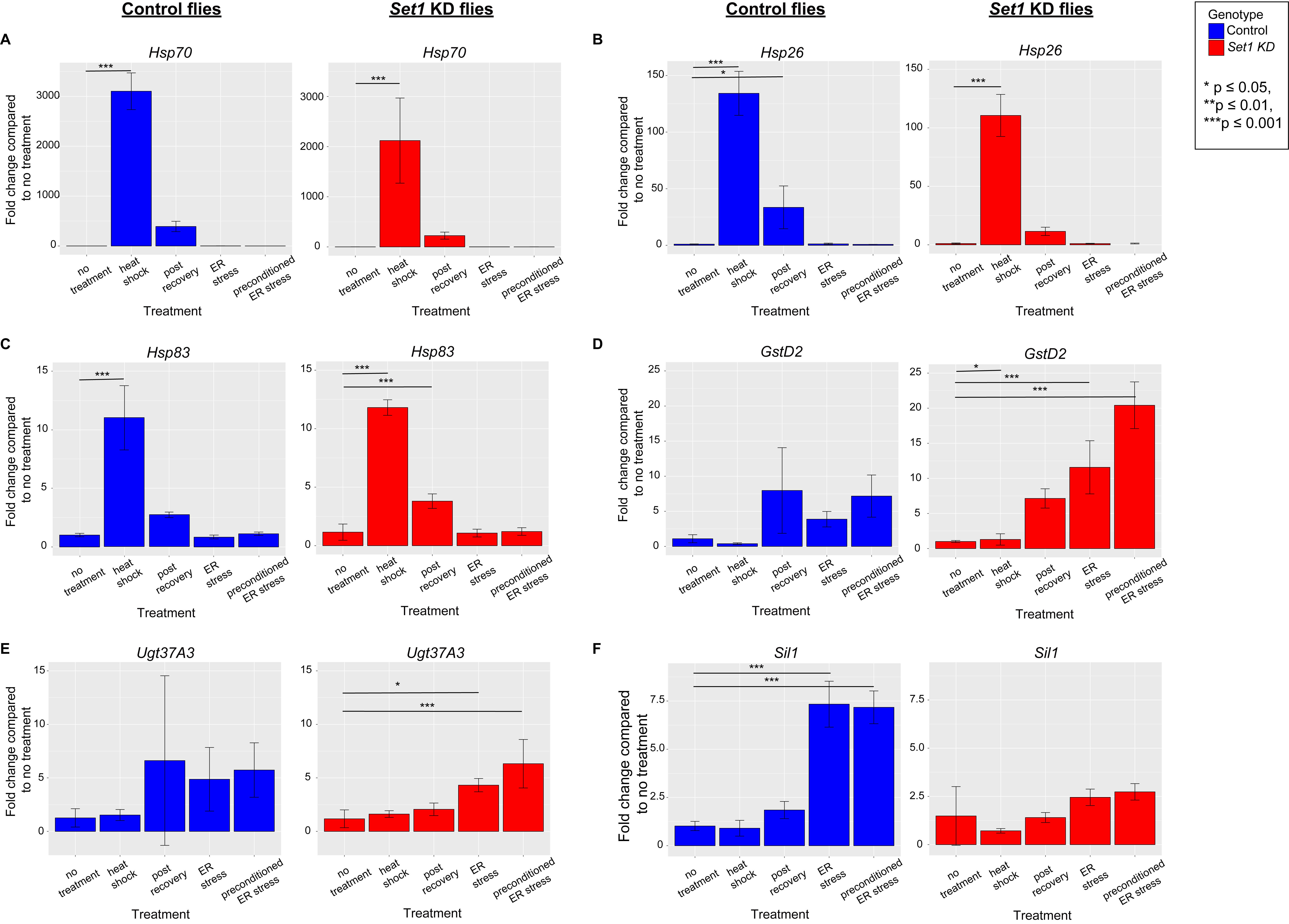
